# Entraining alpha activity using visual stimulation in patients with chronic musculoskeletal pain. A feasibility study

**DOI:** 10.1101/2020.04.27.063339

**Authors:** Laura J. Arendsen, James Henshaw, Christopher A. Brown, Manoj Sivan, Jason R. Taylor, Nelson J. Trujillo-Barreto, Alexander J. Casson, Anthony K. P. Jones

## Abstract

Entraining alpha activity with rhythmic visual, auditory, and electrical stimulation can reduce experimentally induced pain. However, evidence for alpha entrainment and pain reduction in patients with chronic pain is limited. This feasibility study investigated whether visual alpha stimulation can increase alpha power in patients with chronic musculoskeletal pain and secondarily, if chronic pain was reduced following stimulation. In a within-subject design, 22 patients underwent 4-minute periods of stimulation at 10 Hz (alpha), 7 Hz (high-theta, control), and 1 Hz (control) in a pseudo-randomized order. Patients underwent stimulation both sitting and standing and verbally rated their pain before and after each stimulation block on a 0-10 numerical rating scale. Global alpha power was significantly higher during 10 Hz compared to 1 Hz stimulation when patients were standing (t = −6.08, p <.001). On a more regional level, a significant increase of alpha power was found in the right-middle and left-posterior region when patients were sitting. With respect to our secondary aim, no significant reduction of pain intensity and unpleasantness was found. However, only the alpha stimulation resulted in a minimal clinically important difference in at least 50% of participants for pain intensity (50%) and unpleasantness ratings (65%) in the sitting condition. This study provides initial evidence for the potential of visual stimulation as a means to enhance alpha activity in patients with chronic musculoskeletal pain. The brief period of stimulation was insufficient to reduce chronic pain. This study is the first to provide evidence that a brief period of visual stimulation at alpha frequency can significantly increase alpha power in patients with chronic musculoskeletal pain. Further study is warranted to investigate optimal dose and individual stimulation parameters to achieve pain relief in these patients.

## 1 Introduction

Chronic pain is a prevalent and debilitating condition that has a wide-reaching impact on physical and mental well-being (Breivik et al. 2006; Van Hecke, Torrance, and Smith 2013). Opioids and other medications are commonly prescribed to treat chronic pain (Turk, Wilson, and Cahana 2011). However, most medications have considerable side-effects and evidence for their long-term effectiveness is limited (Chou et al. 2015; Turk 2002). In Europe, 40% of people with chronic pain report that their pain was inadequately managed (Breivik et al. 2006). Therefore, the development of alternative therapies to relieve pain is warranted.

Chronic musculoskeletal pain is associated with changes in brain structure and function (Baliki et al. 2014; Brown, El-Deredy, and Jones 2014; Gwilym et al. 2010; Kulkarni et al. 2007). Moreover, chronic pain is associated with changes in oscillatory neural activity. Most commonly, an increase of theta power and beta power has been found (Ploner, Sorg, and Gross 2016; Lim et al. 2016; Sarnthein et al. 2006) and a slowing of the peak alpha frequency (Boord et al. 2008; De Vries et al. 2013; Sarnthein et al. 2006; Lim et al. 2016). Thus, the brain’s response to pain provides a promising target for the development of novel pain therapies (Jensen et al. 2008).

A brain signal of particular interest as a therapeutic target is alpha activity, oscillatory neural activity in the frequency range of 8-12 Hz. Alpha activity gates the processing of incoming sensory information via a mechanism of functional inhibition (Jensen and Mazaheri 2010). Incoming information is gated via the inhibition of brain regions processing irrelevant information (high alpha power), which routes the processing of information to task-relevant regions (low alpha power). This mechanism has been linked to top-down control and attention (Foxe and Snyder 2011; Klimesch 2012) and is also involved in pain processing. Somatosensory alpha activity during pain (Hauck et al. 2015) and the anticipation of pain (May et al. 2012) is modulated by attention, and frontal alpha activity is increased following a placebo-induced expectation of pain relief (Huneke et al. 2013). Importantly, pre-stimulus somatosensory alpha power is inversely related to perceived pain: higher alpha power is associated with lower pain intensity and vice versa, both for experimental pain (Tu et al. 2016; Babiloni et al. 2006) and chronic pain (Camfferman et al. 2017; Ahn et al. 2019). Thus, neurotherapies that increase alpha power may have potential in reducing chronic pain.

Alpha activity can be enhanced through the application of rhythmic stimulation, including visual, auditory, and electrical stimulation (Thut, Schyns, and Gross 2011). When presented with an external stimulation at a certain frequency, oscillatory neural activity at this same frequency synchronizes in phase with the external stimulation, a phenomenon often referred to as entrainment. This ultimately leads to an increase of power at the stimulation frequency at the population level (de Graaf et al. 2013; Spaak, de Lange, and Jensen 2014; Helfrich et al. 2014; Vossen, Gross, and Thut 2015).

Alpha entrainment has been successfully implemented to reduce experimentally induced pain using rhythmic visual (Ecsy, Brown, and Jones 2018), auditory (Ecsy, Jones, and Brown 2017), and transcranial alternating current stimulation (tACS) (Arendsen, Hugh-Jones, and Lloyd 2018). However, to date, only one study successfully induced an increase in somatosensory alpha power that was correlated with a reduction in pain intensity in patients with chronic low-back pain (CLBP), using tACS at alpha frequency over somatosensory regions (Ahn et al. 2019). More work is required to understand the efficacy of alpha entrainment and the relationship between entrainment and analgesia, across different clinical pain populations and using different stimulation modalities.

This feasibility study primarily investigated whether visual alpha stimulation increased global alpha power in patients with chronic musculoskeletal pain, both in a more uncomfortable condition (standing) and a resting condition (sitting). A secondary aim was to explore whether a brief period of alpha stimulation was also associated with reduced clinical pain. Using a within-subject design, alpha stimulation (10 Hz) was compared to 1 Hz control stimulation. Finally, we also compared 7 Hz stimulation (high theta) to 1 Hz stimulation. An important confound to take into account when using visual stimulation to entrain alpha activity, is that synchronization of alpha oscillations can also be induced indirectly via the engagement of attentional mechanisms by the visual stimulus (Thut, Schyns, and Gross 2011; Klimesch 2012; Brüers and Vanrullen 2018). Whereas 7 Hz visual stimulation could lead to indirect synchronization of alpha activity via attentional engagement similar to 10 Hz stimulation, it should not lead to direct alpha entrainment. Thus, the 7 Hz stimulation was included to address the confound of attentional mechanisms related to the rhythmic visual stimulation.

## 2 Materials and methods

### 2.1 Participants

Twenty-two participants were recruited from local pain and musculoskeletal clinics (Salford Royal NHS Trust and North West CATS NHS) and support groups and from the University of Manchester. All participants gave written informed consent to take part in the study and received a reimbursement for their time and travel expenses. The study was approved by the North West - Liverpool East Research Ethics Committee (NHS Health Research Authority; reference number 17/NW/0255).

All participants had a diagnosis of chronic musculoskeletal pain, i.e., pain present for at least 3 months. The study included: 14 patients with fibromyalgia; 2 patients with osteoarthritis; 1 patient with fibromyalgia and osteoarthritis; 2 patients with low back pain; 1 patient with stenosis of the lower back; and 2 patients with widespread chronic pain (no specific diagnosis). All participants took part in a telephone-interview to complete a screening questionnaire prior to participation to ensure they: 1) were aged 18 or older; 2) did not have any difficulty understanding verbal or written English; 3) were not involved in any clinical trials at the time of testing; 4) were not hospitalized/scheduled to be hospitalized during their participation in the study. To ensure it was safe to undergo the rhythmic visual stimulation participants were excluded if they: 1) were diagnosed with epilepsy or had ever had a convulsion or seizure; 2) had any first-degree relative with epilepsy; or 3) had ever experienced discomfort when exposed to flashing lights.

The datasets of two participants were removed from the final analysis as they were not able to complete the entire study due to high levels of pain and discomfort. This resulted in a total of 20 participants that were included in the statistical analysis (mean age ± SD = 43.45 ± 16.82 years; 13 female).

### 2.2 Visual stimulation

All participants underwent visual stimulation at 10 Hz to entrain alpha activity and at the two control frequencies of 1 and 7 Hz. The visual stimulation was administered using goggles with 8 LEDs, 4 around each eye (bespoke equipment – made by Medical Physics, Salford Royal NHS Foundation Trust; Figure 1). Rhythmic flashes were generated using bespoke software run in Matlab 2017a (The Mathworks, Inc., Natick, MA; Matlab). Participants were asked to close their eyes during the stimulation and brightness was adjusted for each individual participant to ensure that stimulation was administered at a comfortable brightness. After participants closed their eyes, they were presented with a 30-second 1 Hz stimulation sample to identify the brightness at which they could clearly perceive the visual stimulation without experiencing any discomfort.

**Figure 1.**
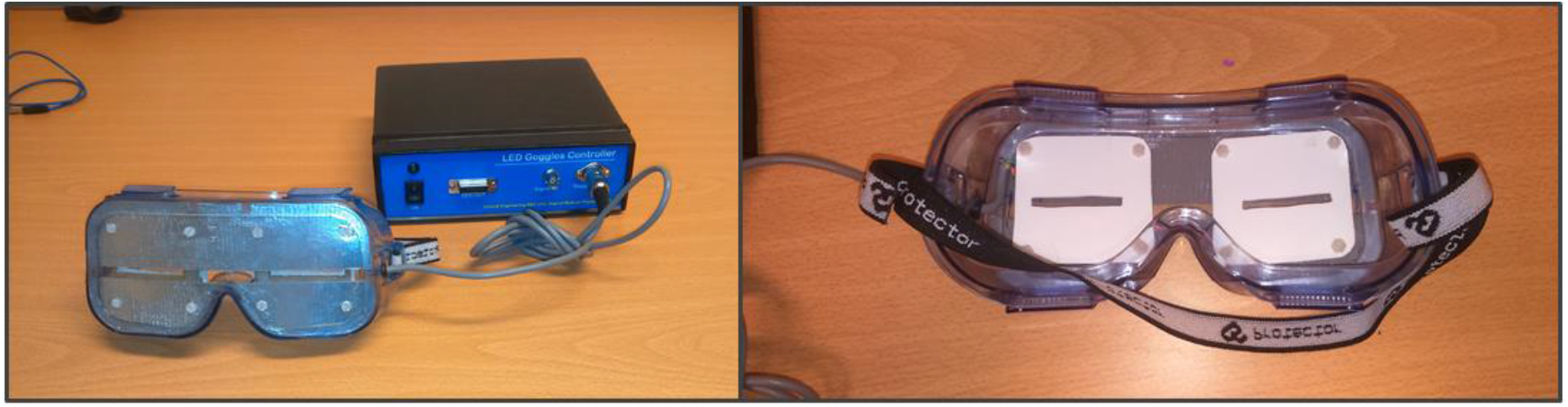
Goggles used for the visual stimulation at the three different stimulation frequencies (bespoke equipment - Medical Physics, Salford Royal NHS Foundation Trust). Eight LEDs were used in total, four around each eye. The goggles were kept in place with an elasticated headband.

The three different stimulation frequencies were delivered in separate blocks (Figure 2). During each block a 1-minute baseline period was followed by 4 minutes of rhythmic visual stimulation, whilst EEG was recorded. During the 1-minute baseline, non-rhythmic visual stimulation was applied with a jittered interstimulus interval (ISI) between flashes. During this non-rhythmic stimulation period the signal phase would change by 180 degrees frequently in a semi-random manner, as not to cause any long-term entrainment effects. These phase changes would occur either every 1.6 s (50% of the time), every 1.15 s (25% of the time), or every 1.9 s (25% of the time). These non-rhythmic baseline periods before each stimulation period were later used for the EEG analysis to provide a standardized baseline for each stimulation condition. It was decided to include the non-rhythmic visual stimulation during the baseline period to ensure that the baseline and entrainment period were kept as similar as possible, e.g. with respect to luminance, with the only difference being that the stimulation during the baseline period was not rhythmic and would therefore not induce any direct alpha entrainment.

**Figure 2.**
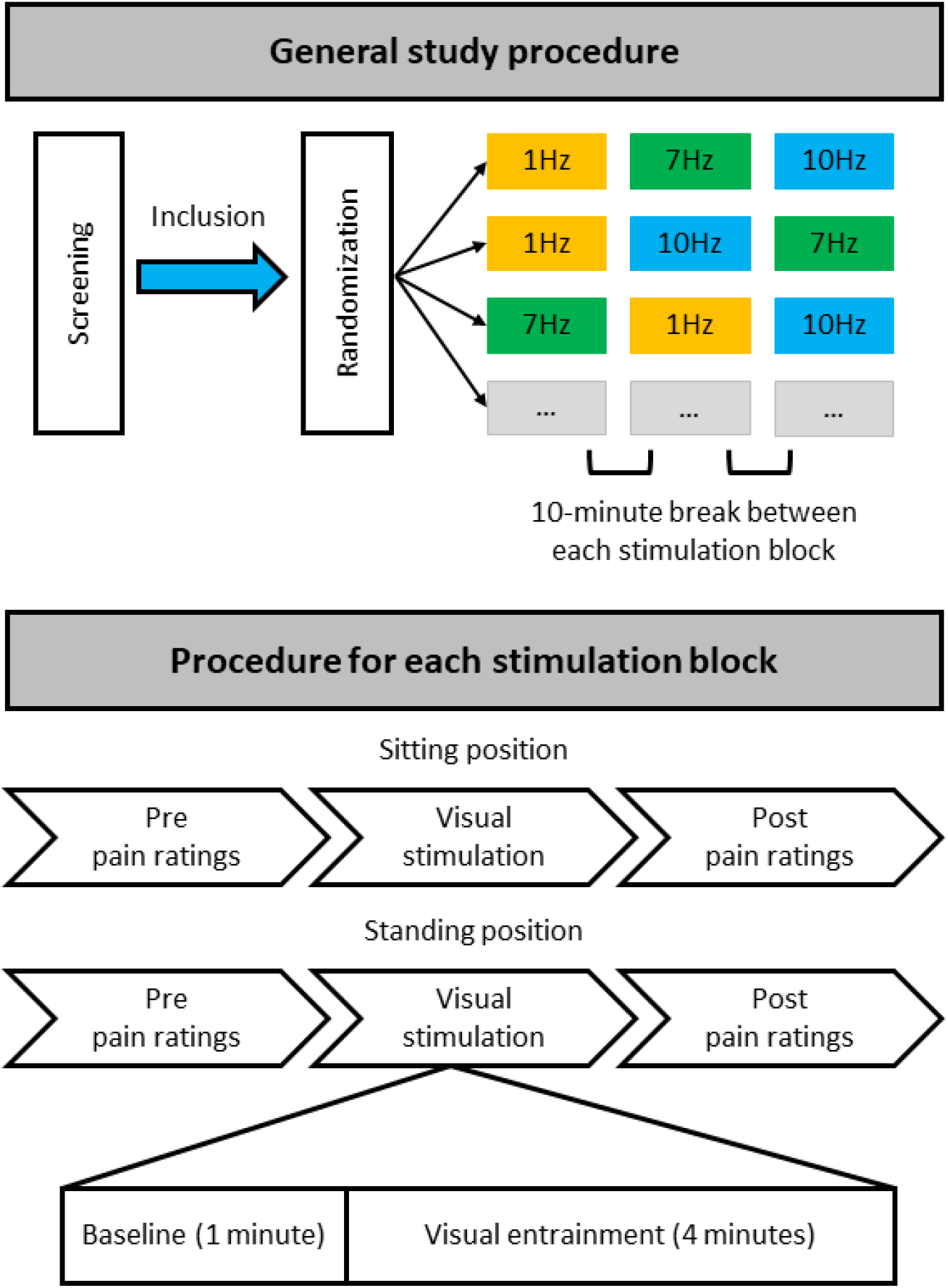
Overview of study procedure. During a single study visit each participant completed three stimulation blocks that each contained stimulation at one particular frequency (1, 7, or 10 Hz), with stimulation both in a sitting and standing position. For each stimulation block half of the participants always started with stimulation whilst sitting, the other half whilst standing. Each participant was randomly allocated to one of six possible stimulation frequency orders: 1, 7, 10 Hz; 1, 10, 7 Hz; 7, 1, 10 Hz; 7, 10, 1 Hz; 10, 1, 7 Hz; or 10, 7, 1 Hz. There was a break of at least 10 minutes between each stimulation block with a specific stimulation frequency, to minimize any potential carry-over effects. Thus, each participant received equal numbers of stimulation blocks at the different frequencies but randomized to a different order of stimulation blocks.

### 2.3 Pain assessment

To quantify chronic pain experience, participants were asked to verbally rate their pain intensity and pain unpleasantness using two 11-point numerical rating scales (NRS) ranging from 0 to 10 (0 = not at all intense/unpleasant, 10 = extremely intense/unpleasant).

To assess the effect of visual alpha stimulation on chronic pain, changes in chronic pain intensity and unpleasantness ratings were assessed in two settings associated with different levels of discomfort: 1) when participants were seated in a comfortable chair (sitting position); and 2) when participants were standing whilst holding the back of the chair for support (standing position). Participants were asked to rate their subjective level of discomfort as a result of the position they were in at the start and end of each stimulation block on an NRS ranging from 0-10 (0 = not at all uncomfortable, 10 = extremely uncomfortable). On average (mean ± SD) patients rated their discomfort at the start of each stimulation block as 3.94 ± 2.14 when sitting and as 4.78 ± 2.07 when standing. At the end of each stimulation block average discomfort was 3.80 ± 2.34 when sitting and 5.53 ± 2.30 when standing. Stimulation was applied both in a setting of higher and lower discomfort to assess whether the level of ongoing discomfort might influence the effect of the visual stimulation.

Pain ratings were collected before and directly after each visual stimulation block, each block including a 1-minute baseline and a 4-minute entrainment period (at 1, 7, and 10 Hz), both in the sitting and standing condition (Figure 2).

### 2.4 Questionnaires

Chronic pain and the outcome of chronic pain treatment is influenced by personality factors and pain-related cognitions and beliefs (Crombez et al. 1999; Granot and Ferber 2005; Keefe et al. 1989). Therefore, a series of questionnaires were included in this study to assess if these questionnaire variables were related to the effect of the visual alpha stimulation. This could potentially inform the design of a future larger trial, e.g., whether to balance participants for these variables between treatment and control arms.

Participants were asked to complete a set of four questionnaires once, during the breaks between stimulation blocks: the Hospital Anxiety and Depression Scale (HADS) (Zigmond and Snaith 1983); the Pain Self-Efficacy Questionnaire (PSEQ) (Nicholas 1989); the Brief Pain Inventory (BPI) (Cleeland and Ryan 1994); and the Multidimensional Health Locus of Control scale (MHLC) (Wallston, Strudler Wallston, and DeVellis 1978).

Both anxiety and depression have been found to frequently co-occur with chronic pain conditions (Mcwilliams, Cox, and Enns 2003). Moreover, a positive association between pain experience and depression and anxiety has been found, both in an experimental pain setting (Tang and Gibson 2005; Walsh 1998) and a clinical pain setting (Geisser et al. 2000; Granot and Ferber 2005). To assess anxiety and depression in the present study we used the HADS. The HADS is a valid self-assessment scale originally developed as a tool to reliably detect states of anxiety and depression in patients attending a general medical clinic (C. Herrmann 1997; Zigmond and Snaith 1983). The HADS comprises of 7 items to assess anxiety and 7 items to assess depression. Participants are asked to tick the box that most closely represents how they were feeling in the past week on a 4-point scale ranging from 0 to 3. For example, “I feel tense or ‘wound up’: (1) not at all; (2) occasionally; (3) a lot of the time; or (4) most of the time.

The BPI, PSEQ and MHLC were used to gain further insight into the pain experience of the participants and their pain-related beliefs and cognitions.

The BPI is a tool to assess both pain intensity (sensory dimension) and pain interference (reactive dimension) in patients with chronic pain. The BPI was originally developed to assess cancer-related pain (Cleeland and Ryan 1994) but is also a widely used and valid measure for patients with non-malignant chronic pain (Tan et al. 2004). Participants are asked to rate their worst and least pain intensity over the last 24 hours, their average pain intensity, and their current pain intensity on a scale of 0 to 10 (sensory dimension). Participants are also asked to rate the degree to which pain interferes with 7 domains of functioning on a scale of 0 to 10, for instance walking ability and relationships with other people (reactive dimension).

The PSEQ is a questionnaire designed to assess self-efficacy beliefs in people experiencing chronic pain (Nicholas 1989), by assessing the confidence participants have in their ability to perform certain tasks and activities despite their pain. The PSEQ contains 10 items describing different settings/activities, such as ‘I can do most of the household chores (e.g. tidying-up, washing dishes, etc.), despite the pain’ and ‘I can live a normal lifestyle, despite the pain’. Participants are asked to rate how confident they are that they can do these things at present despite the pain on a scale from 0 to 6, with 0 = not at all confident and 6 = completely confident.

The MHLC was developed to assess three dimensions of: internal health locus of control, powerful others locus of control, and chance health of control (Wallston, Strudler Wallston, and DeVellis 1978). The MHLC contains 18 items with a belief statement about the participant’s health, for example ‘Whatever goes wrong with my pain condition is my own fault’, and ‘Other people play a big role in whether my pain condition improves, stays the same, or gets worse’. Participants are asked to rate to what extent they agree with each item on a scale of 1 to 6, (1 = strongly disagree and 6 = strongly agree).

### 2.5 EEG acquisition

EEG was recorded during all visual stimulation blocks using 64 Ag/AgCl electrodes attached to a cap according to the extended standard 10-20 system, using the BrainCap MR, BrainAmp DC/MR amplifiers, and the EEG data recording software BrainVision Recorder (Brain Products GmbH, Germany). The FCz electrode was used as a reference electrode and AFz as the ground electrode. EEG was recorded with a sampling rate of 500 Hz and band-pass filter settings of DC-100 Hz.

### 2.6 Procedure

All participants attended the lab for a single study visit during which they underwent visual stimulation at all three frequencies (1, 7, and 10 Hz), once whilst sitting down and once whilst standing up. This resulted in a total of 6 stimulation conditions: three visual stimulation frequencies (1, 7, 10 Hz) x two positions (sitting and standing). For each condition a 1-minute baseline period was followed by a 4-minute stimulation period (Figure 2).

After obtaining written informed consent, completing the EEG setup, and identifying the individual stimulation brightness, each participant was pseudo-randomly allocated to one of the six possible stimulation frequency orders. All participants completed three stimulation blocks, with each block containing one specific stimulation frequency. Each stimulation frequency was experienced both sitting and standing. Half of the participants completed each stimulation block in the sitting position first, the other half started with the standing position first. Pain intensity and unpleasantness were assessed before and after each stimulation condition, i.e., directly before and after the stimulation in the sitting position and also directly before and after the stimulation in the standing position. After each stimulation block participants had a break of at least 10 minutes before carrying on with the next block, to limit potential carry-over effects of the previous block of stimulation. Although both the experimenter and the participants were blinded to the order of visual stimulation frequencies, due to the nature of the stimulation (visual) the frequency of stimulation for each block became apparent as soon as the stimulation was started. However, importantly, participants were not provided with any clues as to which was the expected therapeutic condition.

### 2.7 EEG analysis

The EEG recordings were imported into Matlab (The Mathworks, Inc., Natick, MA; Matlab version R2017a). A number of pre-processing and artifact removal steps were carried out on the continuous EEG data using the EEGLAB toolbox (Delorme and Makeig 2004) in the following order: 1) interpolation of any bad channels (spherical interpolation); 2) re-referencing to the common average; 3) and high-pass (0.05 Hz) and low-pass filtering (30 Hz). A median of 0.5 channels were interpolated with a range from 0 to 5. Next the continuous data was segmented into 2-second consecutive epochs to accommodate later visual inspection of the data post-Independent Component Analysis (ICA). Finally, as EEG for the different stimulation conditions was recorded and saved in separate files, the data from the different stimulation conditions were combined into one single data file per participant. This data was decomposed into independent signals using ICA in order to remove components reflecting artifactual sources, with components from frontal sources reflecting eyeblinks and eye movements selected for removal. The number of ICs to be calculated was adjusted for the number of interpolated channels (N channels – N interpolated channels). The median number of components removed was 2.5 with a range of 1 to 6. The reconstructed EEG data was then visually inspected to remove any remaining muscle artifacts and any other remaining large artifacts, i.e., large spikes and jumps present in the EEG data (on average 4.79% of trials were removed per participant).

Frequency analysis was performed using the Fieldtrip Toolbox (Oostenveld et al. 2011). Average alpha power (8-12 Hz) was calculated for each visual stimulation condition (1, 7, and 10 Hz) using FFT with a single Hanning taper and non-overlapping windows. All individual alpha power outcomes were log-transformed.

Global alpha power was calculated by averaging log-transformed alpha power from 8-12 Hz across all 2-second epochs per condition across all electrodes, resulting in a single average alpha power outcome per visual stimulation condition. The same was applied to the baseline periods preceding each stimulation condition. Next, the average log-transformed alpha power during each stimulation condition was standardized against its respective baseline period (subtraction method: log alpha power entrainment – log alpha power baseline). To assess changes in alpha activity on a more regional level alpha power was also calculated for 9 regions of interest (ROIs) by averaging over the electrodes in each region only (Figure 3).

**Figure 3.**
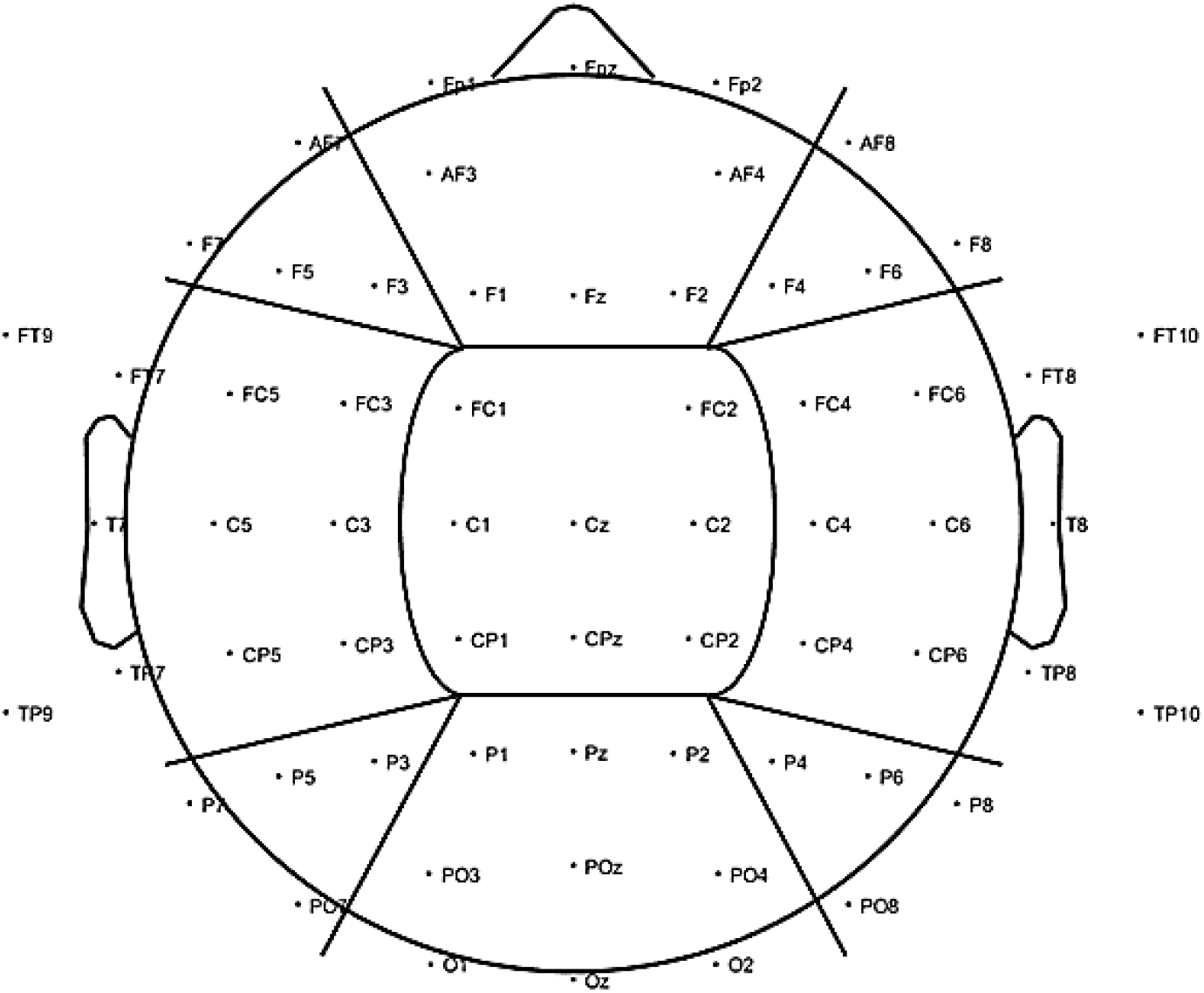
To investigate changes in alpha power (8-12 Hz) for the visual alpha stimulation regionally, further analysis was carried out based on 9 ROIs. We included three anterior ROIs: Left Anterior (LA), including electrodes AF7, F7, F5, and F3; Central Anterior (CA), including electrodes FP1, FPz, FP2, AF3, AF4, F1, Fz, and F2; and Right Anterior (RA): including electrodes AF8, F4, F6, and F8. Three middle ROIs: Left Middle (LM), including electrodes FT9, FT7, FC5, FC3, T7, C5, C3, TP9, TP7, CP5, and CP3; Central Middle (CM), including electrodes FC1, FC2, C1, Cz, C2, CP1, CPz, and CP2; and Right Middle (RM): including electrodes FC4, FC6, FT8, FT10, C4, C6, T8, CP4, CP6, TP8, and TP10. Finally, three posterior ROIs: Left Posterior (LP), including electrodes P7, P5, P3, and PO7; Central Posterior (CP), including electrodes P1, Pz, P2, PO3, POz, PO4, O1, Oz, and O2; and Right Posterior (RP): including electrodes P4, P6, P8, and PO8.

### 2.8 Statistical analysis

Statistical analysis was performed using SPSS version 22 (IBM Corp, Armonk, NY, USA). To assess the effect of the visual stimulation on global alpha power, i.e. alpha power averaged across all electrodes, a repeated-measures ANOVA with the factors Stimulation (1, 7, and 10 Hz) and Position (sitting and standing) was applied.

Next, to further explore the effect of alpha stimulation (10 Hz) on a more regional level, changes in alpha power were also compared for the 9 ROIs (Figure 3). Specifically, 10 Hz stimulation was compared to the 1 Hz control stimulation using a repeated-measures ANOVA with the factors Stimulation (1 and 10 Hz), Position (sitting and standing), the Left-to-Right (L-R) ROI factor (left, central, and right), and the Anterior-to-Posterior (A-P) ROI (anterior, middle, and posterior).

Finally, two repeated-measures ANOVAs with the factors Stimulation (1, 7, and 10 Hz), Position (sitting and standing), and Time (pre- and post-stimulation) were applied to assess a change in pain intensity and unpleasantness ratings respectively. In the case of a violation of sphericity the Greenhouse-Geisser corrected outcomes were used. To correct for multiple comparisons, the Bonferroni correction was applied.

### 2.9 Minimal clinically important difference (MCID) in pain ratings

In line with the recommendations of the Initiative on Methods, Measurement, and Pain Assessment in Clinical Trials (IMMPACT) consensus statement (Dworkin et al. 2005), we also assessed what percentage of participants showed a minimally important clinical difference (MCID) in pain intensity and unpleasantness. A MCID is considered the smallest difference in pain rating that patients perceive as important and is reflected by a 15% reduction in pain intensity/unpleasantness rating compared to baseline ((pain rating post – pain rating pre) / pain rating pre) (Dworkin et al. 2008).

### 2.10 Correlations

To assess the potential relationship between alpha activity and changes in pain due to the alpha stimulation, correlations between standardized global alpha power and the change in pain intensity/unpleasantness rating were calculated. In detail, the average log-transformed alpha power during 10 Hz stimulation standardized against its baseline period was used (subtraction method: alpha power entrainment – alpha power baseline). To calculate the change in the pain intensity/unpleasantness ratings we used: ratings post – ratings pre. Thus, a negative correlation would reflect that higher alpha power during stimulation (compared to baseline) was associated with lower pain ratings post-stimulation and vice versa.

To assess the relationship between changes in pain following alpha stimulation and personality factors and pain-related cognitions and beliefs, correlations between the different questionnaire scores and the change in pain intensity and unpleasantness rating were calculated. The following questionnaire outcomes were used: the sum score for the depression subscale and anxiety subscale separately (HADS); the individual rating for average, worst, and least pain intensity over the last 24 hours and a sum score for the seven pain interference items (BPI); a single sum score for all PSEQ items; and separate sum score for each of the three subscales of the MHLC.

## 3 Results

### 3.1 Global alpha power

The repeated-measures ANOVA with the factors Stimulation (1, 7, and 10 Hz) and Position (sitting and standing) showed a significant main effect of Stimulation (F 2,38 = 34.88; p < .001; Partial Eta^2^ = .65) on global alpha power and a significant interaction between Stimulation and Position (F2,38 = 13.48; p = <.001; Partial Eta^2^ = .42; Figure 4). Post-hoc repeated-measures t-tests showed that alpha power was significantly higher during 10 Hz stimulation compared to the 1 Hz control stimulation during the standing condition (t = −6.08, p <.001). There was no significant increase of global alpha power during the sitting condition (t = −1.30, p = .21). No increase of alpha power was found for the 7 Hz condition compared to 1 Hz condition, only a significant decrease of global alpha power compared to 1 Hz stimulation in the sitting condition (t = 2.51, p = .021). However, this effect did not survive correction of multiple comparisons (corrected significance level of .0125).

**Figure 4.**
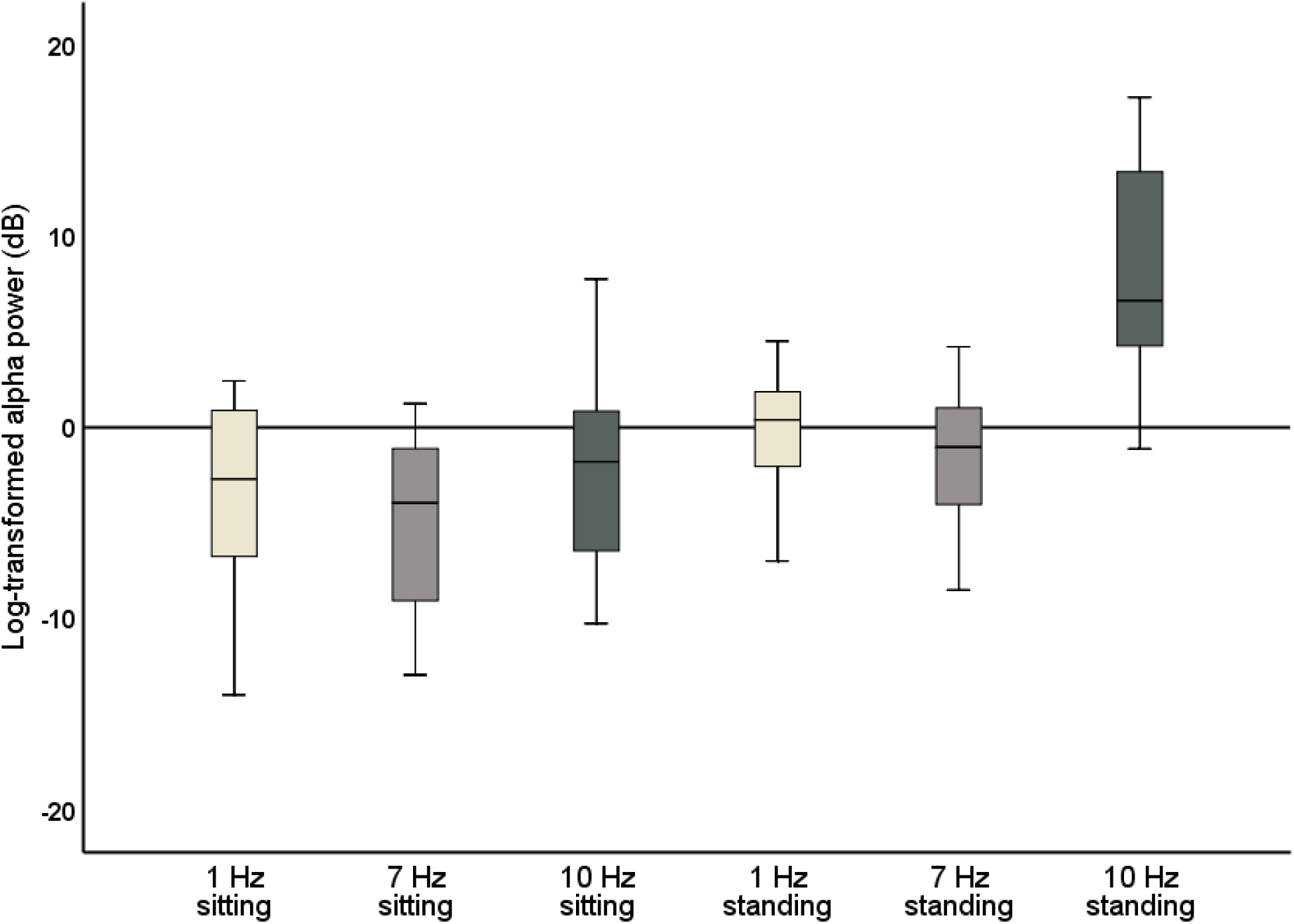
Boxplots of global alpha power (8-12 Hz) during 1 Hz, 7 Hz, and 10 Hz visual stimulation standardized against their respective baseline period, i.e. the change in global alpha power during stimulation compared to baseline.

### 3.2 ROI alpha power analysis

A repeated-measures ANOVA was calculated to assess more regional changes in alpha power with the factors Stimulation (1, 7, and 10 Hz), Position (sitting and standing), L-R ROI (left, central, and right) and A-P ROI (anterior, middle, and posterior). A significant main effect was found for the A-P ROI factor (F_1.34,25.49_ = 11.43; p = .001; Partial Eta^2^ = .38). Moreover, a significant interaction between Stimulation and A-P ROI (F_2.55,48.35_ = 1.73; p = .18; Partial Eta^2^ = .083) was found, and a significant interaction between Stimulation, Position, and A-P ROI (F_2.48,47.07_ = 6.65; p < .002; Partial Eta^2^ = .26). Similarly, a significant main effect was found for the L-R ROI factor (F_1.25,23.76_ = 4.50; p = .037; Partial Eta^2^ = .19), accompanied by a significant interaction between Stimulation and L-R ROI (F_1.25,23.68_ = 11.87; p = .001; Partial Eta^2^ = .39) and a significant interaction between Stimulation, Position, and the L-R ROI (F_1.91,36.20_ = 12.24; p = < .001; Partial Eta^2^ = .39). Thus, the effect of the visual stimulation on alpha power was different depending on position and on scalp region.

Post-hoc repeated-measures t-tests showed that alpha power was significantly higher during 10 Hz stimulation compared to 1 Hz stimulation in the right-middle region (RM) and the left-posterior region (LP) in particular when participants were sitting (Table 1, Figure 5). In the standing condition, alpha power was higher during 10 Hz compared to 1 Hz stimulation across a wider range of mostly middle-anterior regions, however these effects did not survive correction for multiple comparisons (corrected significance level of .0056).

**Figure 5.**
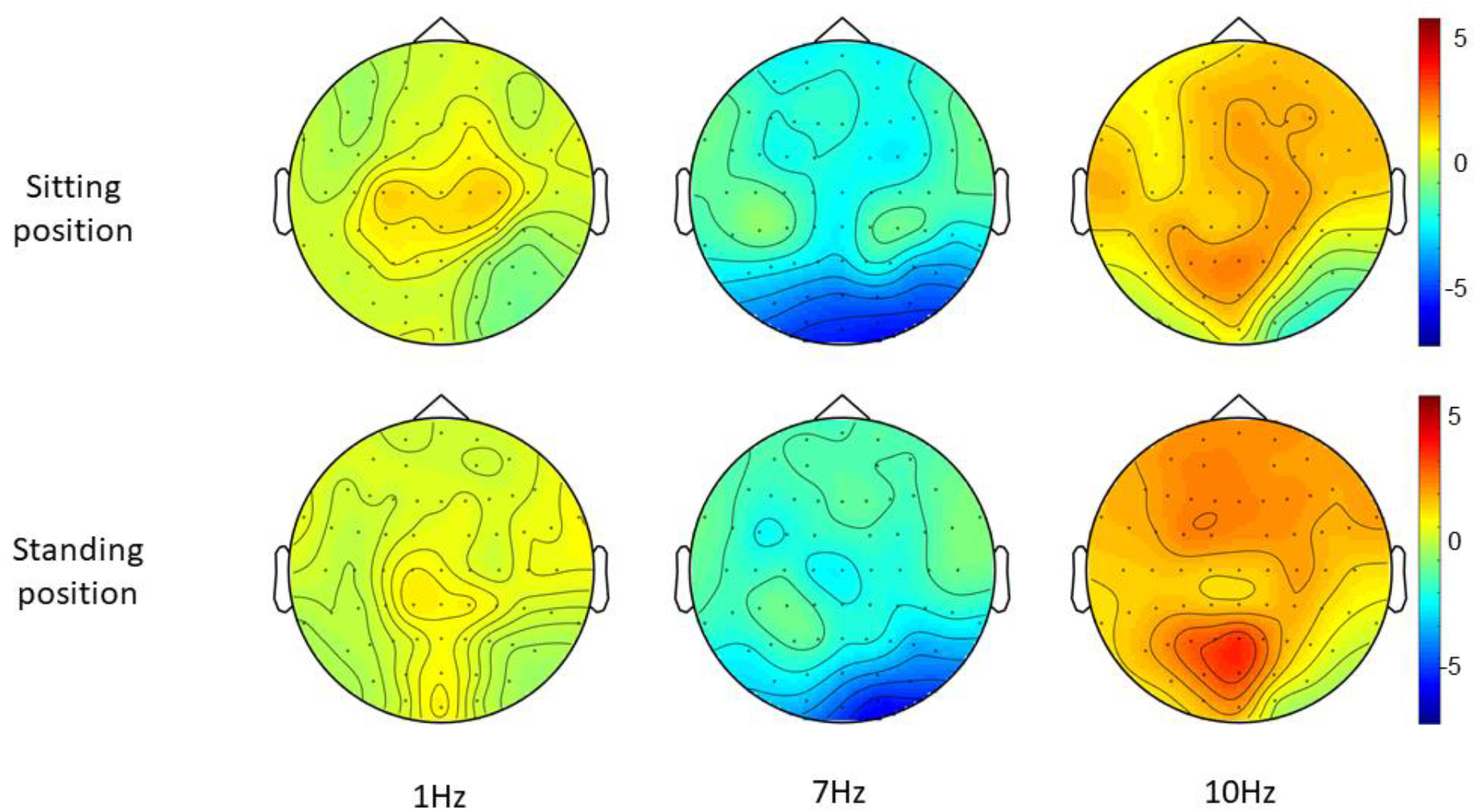
Topographies of standardized alpha power (8-12 Hz), i.e. the change in alpha power during each entrainment condition compared to their respective baseline period.

**Table 1.**
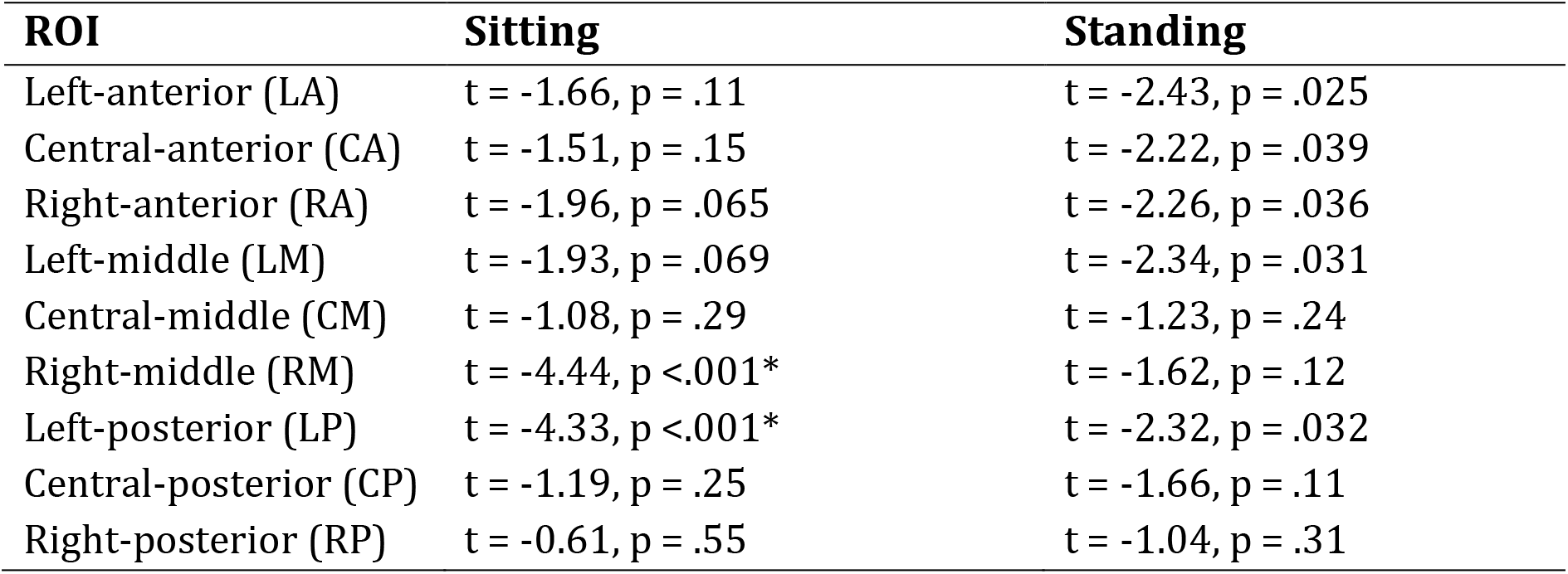
Outcomes of the post-hoc repeated measures t-tests comparing 1 and 10 Hz stimulation for the 9 ROIs separately. Significant t-tests are marked with a * (Bonferroni-corrected significance level).

| ROI | Sitting | Standing |
| --- | --- | --- |
| Left-anterior (LA) | t = -1.66, p = .11 | t = -2.43, p = .025 |
| Central-anterior (CA) | t = -1.51, p = .15 | t = -2.22, p = .039 |
| Right-anterior (RA) | t = -1.96, p = .065 | t = -2.26, p = .036 |
| Left-middle (LM) | t = -1.93, p = .069 | t = -2.34, p = .031 |
| Central-middle (CM) | t = -1.08, p = .29 | t = -1.23, p = .24 |
| Right-middle (RM) | t = -4.44, p < .001* | t = -1.62, p = .12 |
| Left-posterior (LP) | t = -4.33, p < .001* | t = -2.32, p = .032 |
| Central-posterior (CP) | t = -1.19, p = .25 | t = -1.66, p = .11 |
| Right-posterior (RP) | t = -0.61, p = .55 | t = -1.04, p = .31 |

Post-hoc t-tests comparing 1 Hz and 7 Hz visual stimulation showed that alpha power was significantly lower during 7 Hz stimulation compared to 1 Hz stimulation, in particular in the central-middle region (CM) and the central-posterior region (CP; Table 2, Figure 5), when participants were sitting. A similar pattern was present when participants were standing, however this did not survive correction for multiple comparisons for the central-middle region.

**Table 2.** Outcomes of the post-hoc repeated measures t-tests comparing 1 and 7 Hz stimulation for the 9 ROIs. Significant t-tests are marked with a * for the Bonferroni-corrected significance level.

| ROI | Sitting | Standing |
| --- | --- | --- |
| Left-anterior (LA) | t = 2.44, p = .025 | t = 2.01, p = .059 |
| Central-anterior (CA) | t = 3.08, p = .006 | t = 1.73, p = .099 |
| Right-anterior (RA) | t = 2.70, p = .014 | t = 1.93, p = .068 |
| Left-middle (LM) | t = 2.87, p = .010 | t = 2.23, p = .038 |
| Central-middle (CM) | t = 3.66, p = .002* | t = 3.03, p = .007 |
| Right-middle (RM) | t = 2.76, p = .012 | t = 2.57, p = .019 |
| Left-posterior (LP) | t = -1.82, p = .085 | t = 1.99, p = .062 |
| Central-posterior (CP) | t = 4.44, p < .001* | t = 3.45, p = .003* |
| Right-posterior (RP) | t = 2.66, p = .016 | t = 2.73, p = .013 |

### 3.3 Intensity ratings

The repeated-measures ANOVA demonstrated a significant main effect of Position (sitting and standing) on intensity ratings (F_1,19_ = 12.32; p = .002; Partial Eta^2^ = .39), but no significant main effect of Stimulation (F_2,38_ = 1.80; p = .18; Partial Eta^2^ = .087). There was not an interaction between significant Stimulation and Time (pre- and post-stimulation) (F_1.46,27.80_ = .065; p = .89; Partial Eta^2^ = .003), nor a significant interaction between Stimulation, Position, and Time (F_2,38_ = .59; p = .56; Partial Eta^2^ = .030) (Table 3, Figure 6). Moreover, the repeated-measures t-tests further assessing an effect on pain intensity for the alpha stimulation (10 Hz) specifically, did not find a significant change in pain intensity ratings comparing pre- and post-alpha stimulation (sitting: t = 1.54, p = .014; and standing: t = −1.11, p = .28).

**Figure 6.**
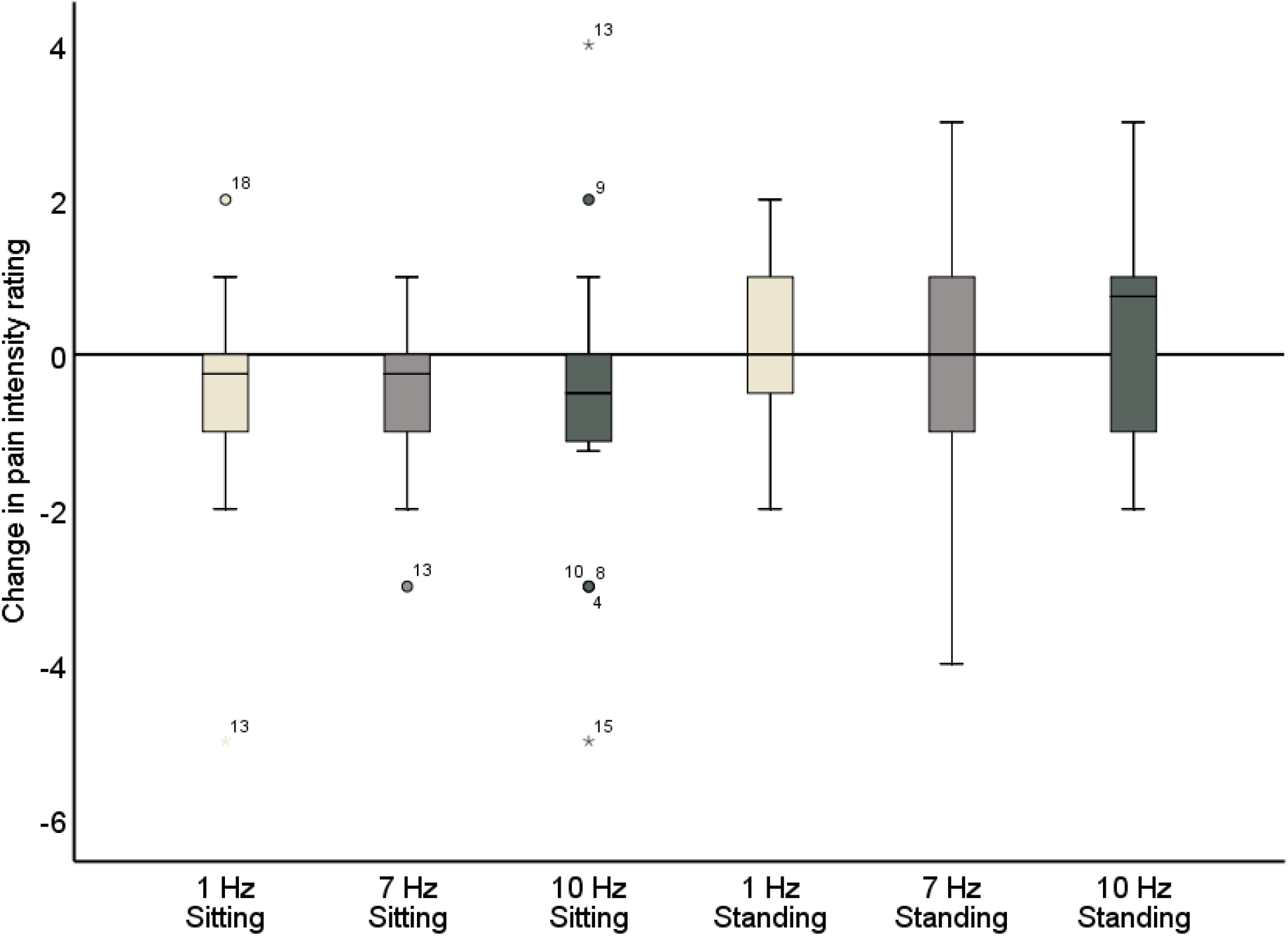
Change in pain intensity ratings comparing pre- and post-stimulation for 1, 7, and 10 Hz stimulation, both in the sitting and standing condition. A negative score reflects a reduction in pain and a positive score reflects an increase of pain following the stimulation.

**Table 3.** Pain intensity ratings (Mean ± SD) pre- and post-stimulation, for the 1, 7, and 10 Hz stimulation condition and for the sitting and standing position.

| Intensity ratings |  |  |  |  |
| --- | --- | --- | --- | --- |
| ROI | Sitting Pre | Sitting Post | Standing Pre | Standing Post |
| 1 Hz | 4.70 ± 1.95 | 4.15 ± 2.20 | 5.18 ± 1.84 | 5.28 ± 1.97 |
| 7 Hz | 4.43 ± 1.70 | 4.03 ± 2.07 | 4.75 ± 1.99 | 4.60 ± 2.47 |
| 10 Hz | 4.39 ± 2.12 | 3.73 ± 2.01 | 4.45 ± 1.87 | 4.78 ± 2.07 |

### 3.4 Unpleasantness ratings

The repeated measures ANOVA demonstrated a significant main effect of Position (sitting and standing) on unpleasantness ratings (F_1,19_ = 12.61; p = .002; Partial Eta^2^ = .40), but no significant main effect of Stimulation (F_2,38_ = 1.78; p = .18; Partial Eta^2^ = .085). There was also not a significant interaction between Stimulation and Time (F_2,38_ = .73; p = .49; Partial Eta^2^ = .037), nor a significant interaction between Stimulation, Position, and Time (F_2,38_ = 2.63; p = .085; Partial Eta^2^ = .12) (Table 4, Figure 7). Moreover, the repeated-measures t-tests assessing an effect on pain unpleasantness for the alpha stimulation (10 Hz) specifically did not find a significant change of pain unpleasantness ratings comparing pre- and post-alpha stimulation (sitting: t = 1.77, p = .093; and standing: t = 1.32, p = .20).

**Figure 7.**
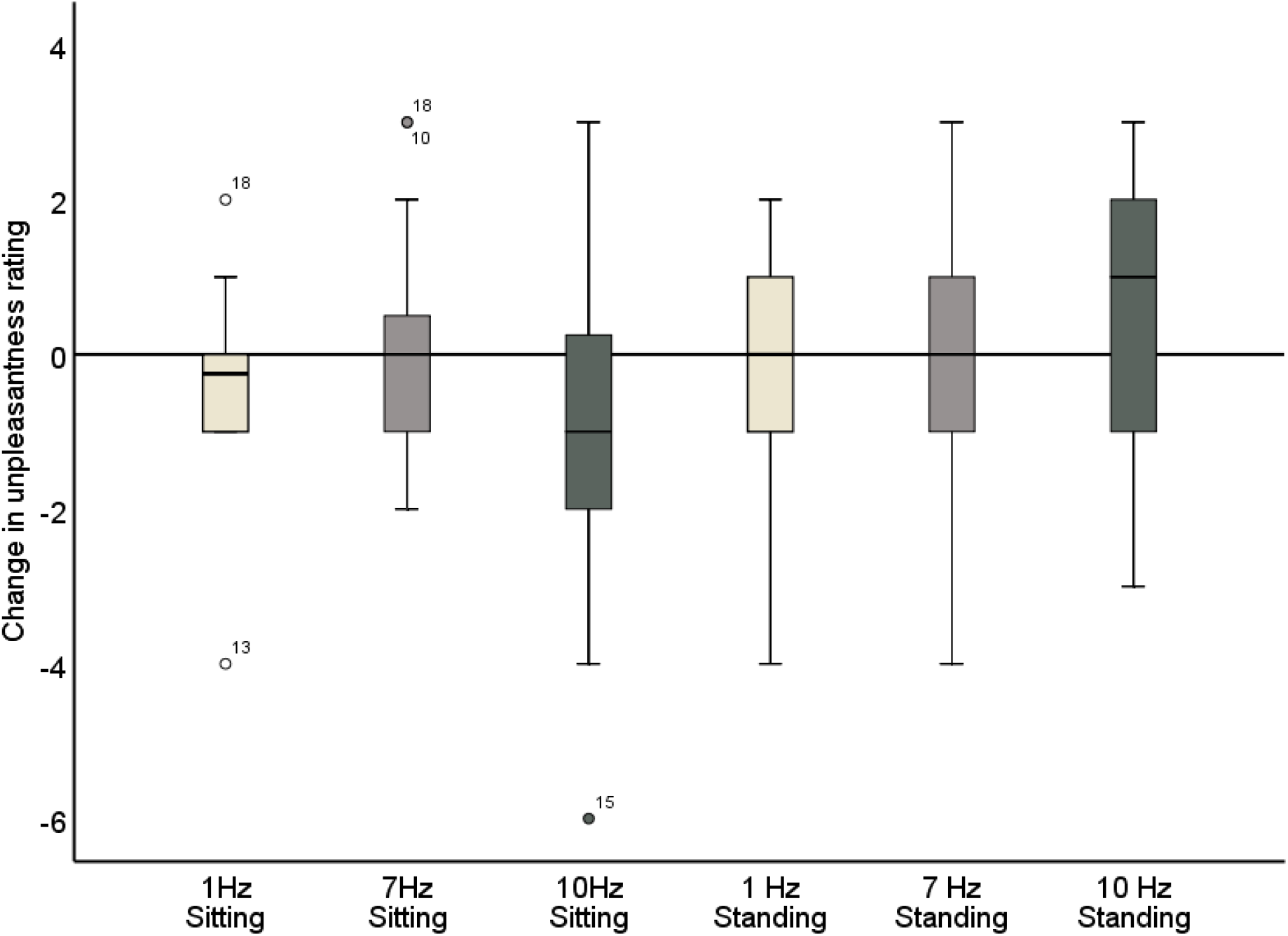
Change in pain unpleasantness ratings comparing pre- and post-stimulation for 1, 7, and 10 Hz stimulation, both in the sitting and standing condition. A negative score reflects a reduction in pain and a positive score reflects an increase of pain following the stimulation.

**Table 4.** Pain unpleasantness ratings (Mean ± SD) pre- and post-stimulation, for the 1, 7, and 10 Hz stimulation condition and for the sitting and standing position.

| Unpleasantness ratings |  |  |  |  |
| --- | --- | --- | --- | --- |
| ROI | Sitting Pre | Sitting Post | Standing Pre | Standing Post |
| 1 Hz | 4.55 ± 2.04 | 4.10 ± 2.37 | 5.40 ± 1.98 | 5.20 ± 2.02 |
| 7 Hz | 4.20 ± 1.82 | 4.33 ± 2.34 | 4.63 ± 1.75 | 4.65 ± 2.24 |
| 10 Hz | 4.38 ± 2.31 | 3.50 ± 2.07 | 4.28 ± 1.89 | 4.80 ± 2.17 |

### 3.5 Minimal clinically important difference (MCID) in pain ratings

#### Sitting condition

We assessed the number of participants that showed a MCID (percentage change > 15%) in pain ratings, for the three different stimulation conditions separately. For the intensity ratings, 50% of participants demonstrated a MCID for the 10 Hz stimulation. A similar value was found for the 1 Hz condition (45%). For the 7 Hz condition a MCID was found for 35%. For the unpleasantness ratings, 65% of participants demonstrated a MCID for the 10 Hz stimulation. This percentage was lower for the 1 Hz condition (40%) and the 7 Hz condition (30%). Only the alpha stimulation led to a MCID in pain intensity and unpleasantness for ≥ 50% of participants.

To further explore any potential pattern of responsiveness, e.g., whether the participants tended to either improve for all stimulation frequencies or not at all, or whether participants improved for only one particular stimulation frequency, an overview of MCIDs for individual participants for each stimulation frequency was created for pain intensity (Table 5) and pain unpleasantness (Table 6).

**Table 5.** All individual participants’ responses to the three stimulation conditions when they were sitting, showing whether they had a MCID for each respective stimulation frequency or no MCID in pain intensity ratings for any of the stimulation conditions. Patient IDs with a single * represent participants that had a MCID of pain intensity for two out of three stimulation conditions and two ** represent participants that had a MCID for all three the stimulation conditions.

|  | N | Patient ID number |
| --- | --- | --- |
| No MCID for all stimulation freq. | 5 | 136, 138, 140, 153, 155 |
| 1 Hz MCID | 9 | 137, 144, 145*, 146**, 148*, 150**, 151, 152*, 154* |
| 7 Hz MCID | 7 | 139*, 141*, 143*, 146, 148, 149, 150 |
| 10 Hz MCID | 10 | 139, 141, 142, 143, 145, 146, 147, 150, 152, 154 |

**Table 6.** All individual participants’ responses to the three stimulation conditions when they were sitting, showing whether they had a MCID for each respective stimulation frequency or no MCID in pain unpleasantness ratings for any of the stimulation conditions. Patient IDs with a single * represent participants that had a MCID of pain unpleasantness for two out of three stimulation conditions and two ** represent participants that had a MCID for all three the stimulation conditions.

|  | N | Patient ID number |
| --- | --- | --- |
| No MCID for all stimulation freq. | 3 | 138, 140, 153 |
| 1 Hz | 8 | 137, 144, 146**, 148, 150**, 151, 152**, 154* |
| 7 Hz | 6 | 139*, 143*, 146, 149*, 150, 152 |
| 10 Hz | 13 | 136, 139, 141, 142, 143, 145, 146, 147, 149, 150, 152, 154, 155 |

From the 20 participants, 6 participants demonstrated an MCID of intensity ratings for 1 stimulation frequency only, 7 participants for two stimulation frequencies, and 2 participants for all three stimulation frequencies. Five participants did not show a MCID in pain intensity for any stimulation condition. For the unpleasantness ratings, 10 participants demonstrated an MCID for 1 stimulation frequency only, 4 participants for two stimulation frequencies, and 3 participants for all three stimulation frequencies. Three participants did not show a MCID of unpleasantness ratings for any stimulation condition. To conclude, there was no clear pattern in responsiveness present when participants were sitting, within the group of participants the change in pain intensity and unpleasantness across the three different stimulation conditions varied considerably.

#### Standing condition

Next, we assessed the number of participants that showed a MCID (percentage change > 15%) in pain ratings, for the three different stimulation conditions separately when the participants were standing. For the intensity ratings, 25% of participants demonstrated a MCID (percentage change > 15%) for the 10 Hz stimulation. A similar value was found for the 1 Hz condition (20%) and the 7 Hz condition (30 %). For the unpleasantness ratings, 30% of participants demonstrated a MCID for the 10 Hz stimulation. This percentage was also found for the 1 Hz condition (30%) and the 7 Hz condition (30%). None of the stimulation conditions led to a MCID in pain intensity or unpleasantness for ≥ 50% of participants in the standing condition.

Next, any potential pattern of responsiveness was also assessed for the standing condition, for pain intensity (Table 7) and pain unpleasantness (Table 8). From the 20 participants, 6 participants demonstrated an MCID of intensity ratings for 1 stimulation frequency only, 3 participants for two stimulation frequencies, and 1 participant for all three stimulation frequencies. Ten participants did not show a MCID in pain intensity for any stimulation condition. For the unpleasantness ratings, 5 participants demonstrated an MCID for 1 stimulation frequency only, 5 participants for two stimulation frequencies, and 1 participant for all three stimulation frequencies. Nine participants did not show a MCID of unpleasantness ratings for any stimulation condition. To conclude, there was also no clear pattern in responsiveness present when the participants were standing. Moreover, in the standing condition there was a larger group of participants that did not demonstrate a MCID for any of the stimulation conditions compared to the sitting condition.

**Table 7.** All individual participants’ responses to the three stimulation conditions when they were standing, showing whether they had a MCID for each respective stimulation frequency or no MCID in pain intensity ratings for any of the stimulation conditions. Patient IDs with a single * represent participants that had a MCID of pain intensity for two out of three stimulation conditions and two ** represent participants that had a MCID for all three the stimulation conditions.

|  | N | Patient ID number |
| --- | --- | --- |
| No MCID for all stimulation freq. | 10 | 136, 138, 139, 140, 142, 146, 147, 151, 153, 155 |
| 1 Hz MCID | 4 | 141*, 144*, 145**, 150* |
| 7 Hz MCID | 6 | 143, 144, 145, 148, 150, 152 |
| 10 Hz MCID | 5 | 137, 141, 145, 149, 154 |

**Table 8.**
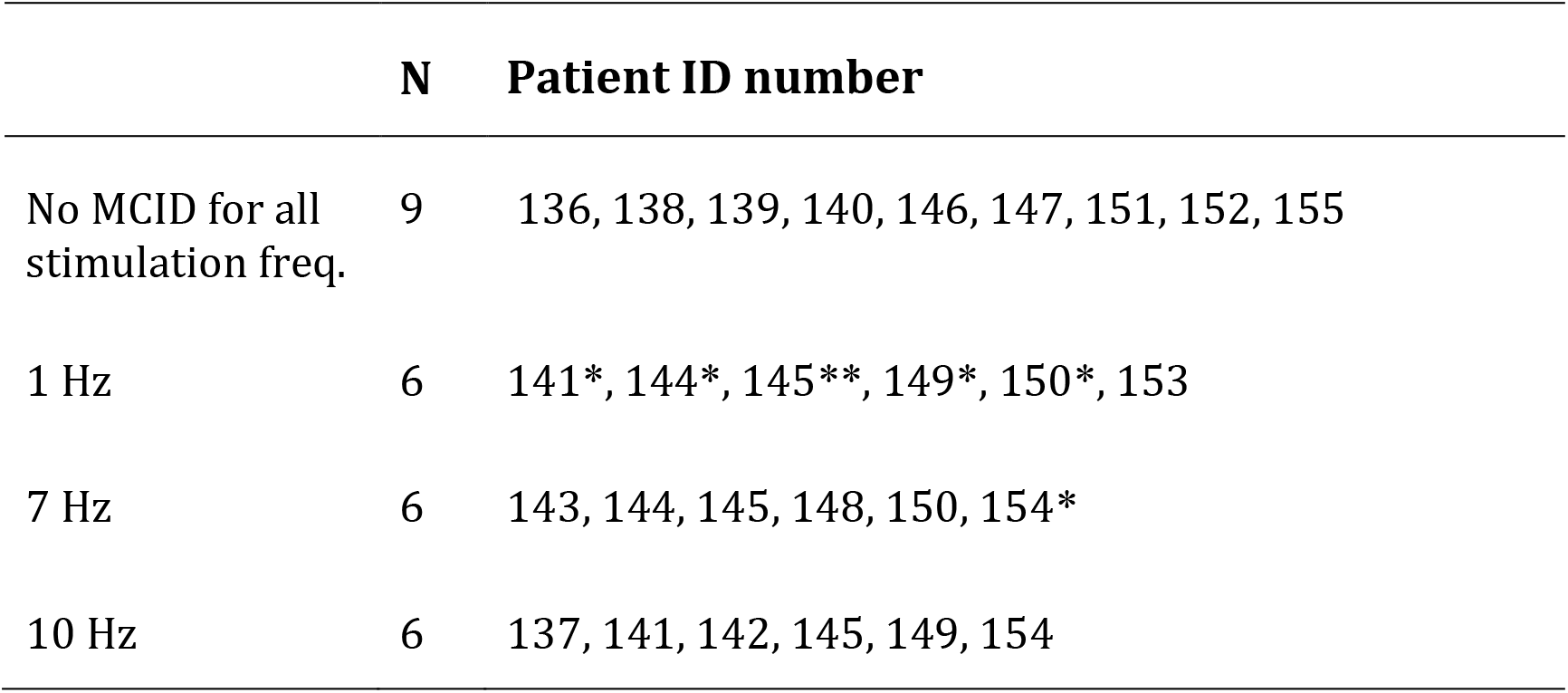
All individual participants’ responses to the three stimulation conditions when they were standing, showing whether they had a MCID for each respective stimulation frequency or no MCID in pain unpleasantness ratings for any of the stimulation conditions. Patient IDs with a single * represent participants that had a MCID of pain unpleasantness for two out of three stimulation conditions and two ** represent participants that had a MCID for all three the stimulation conditions.

|  | N | Patient ID number |
| --- | --- | --- |
| No MCID for all stimulation freq. | 9 | 136, 138, 139, 140, 146, 147, 151, 152, 155 |
| 1 Hz | 6 | 141*, 144*, 145**, 149*, 150*, 153 |
| 7 Hz | 6 | 143, 144, 145, 148, 150, 154* |
| 10 Hz | 6 | 137, 141, 142, 145, 149, 154 |

### 3.6 Correlations

The correlations between standardized global alpha power during 10 Hz stimulation (log alpha power during 10 Hz stimulation – baseline log alpha power) and the difference in intensity/unpleasantness ratings comparing pre- and post-stimulation were calculated (ratings post – ratings pre). No significant correlation was found for the intensity ratings (Sitting: r = .34; p = .14; N = 20. Standing: r = .16; p = .51; N = 20). For the unpleasantness ratings no significant correlation with global alpha power was found either for the standing condition (r = .11; p = .65; N = 20). A correlation was found for the sitting condition (r = .46; p = .04; N = 20), however this did not survive correction for multiple comparisons (corrected significance level = .0125).

Post hoc it was decided to also explore the correlations between the change in pain intensity/unpleasantness ratings and standardized alpha power for the two ROIs that showed a significant increase of alpha power during 10 Hz stimulation compared to 1 Hz stimulation in the sitting condition, the right-middle (RM) and left-posterior (LP) ROI. For the RM ROI, there was no significant correlation between change in pain ratings (ratings post – ratings pre) and alpha power (intensity ratings: r = -.40; p = .082; N = 20; unpleasantness ratings: r = -.41; p = .073; N = 20). For the LP ROI, no significant correlation was found either for the intensity ratings (r = .43; p = .060; N = 20). A correlation was identified for the unpleasantness ratings (r = .54; p = .015; N = 20), but this did not survive correction for multiple comparisons (corrected significance level = .0125).

Finally, correlations were assessed between the change in intensity/unpleasantness ratings and the questionnaire outcomes. No significant correlation between ratings and any of the questionnaire outcomes was found. A correlation between pain unpleasantness ratings and the HADS Depression subscale was identified in the standing condition (r = .50; p = .026; N = 20). However, this did not survive correction for multiple comparisons.

## 4 Discussion

Emerging evidence shows an inverse relationship between alpha power and chronic pain (Camfferman et al. 2017; Ahn et al. 2019). Therefore, alpha activity has been proposed as a key target for novel neuromodulatory therapies to manage chronic pain (M. P. Jensen et al. 2008). This feasibility study primarily aimed to assess the efficacy of visual alpha stimulation to enhance alpha activity in patients with chronic musculoskeletal pain. Secondarily, it was evaluated whether a brief period of alpha stimulation was also sufficient to reduce chronic pain. The main finding of this study was that visual alpha stimulation can effectively enhance alpha activity in patients with chronic musculoskeletal pain. Global alpha power was significantly higher during alpha stimulation compared to the 1 Hz control stimulation when patients were experiencing stronger discomfort (standing condition). On a more regional level, a significant increase of alpha activity was also found in the right-middle and left-posterior region when patients were sitting. With respect to our secondary aim, four minutes of alpha stimulation was not sufficient to significantly reduce chronic pain. However, only the alpha stimulation resulted in a MCID in at least 50% of participants for the pain intensity (50%) and unpleasantness ratings (65%). This study is the first to demonstrate the efficacy of rhythmic visual stimulation to modulate alpha activity in patients with chronic pain. However, further study is warranted to investigate the optimal dose and individual stimulation parameters (Krause and Cohen Kadosh 2014) such as duration and frequency of entrainment to achieve significant pain relief.

Whereas both 7 and 10 Hz stimulation can result in an indirect entrainment of alpha activity via attentional mechanisms (Thut, Schyns, and Gross 2011), only the 10 Hz stimulation should lead to a direct entrainment of alpha. No increase of alpha power was found for the 7 Hz stimulation compared to 1 Hz stimulation. This suggests that the effect of alpha stimulation on alpha power found in this study is likely the result of direct entrainment, and not only reflecting a non-specific effect of attention being directed away from the pain by the visual stimulation.

The present study’s findings build on the findings by Ecsy et al.(Ecsy, Brown, and Jones 2018), who previously demonstrated that visual alpha stimulation can increase alpha power and reduce pain in an experimental pain setting. Ahn et al. (2019) provided the first evidence that alpha stimulation can be used successfully in a clinical pain setting. They demonstrated that alpha tACS applied over somatosensory regions enhances somatosensory alpha power in patients with CLBP. Here we demonstrate that rhythmic visual stimulation can also modulate alpha activity in patients. Moreover, as this study included patients with various chronic musculoskeletal pain conditions, it also offers a first indication that the modulation of alpha activity with alpha stimulation can be generalized across different chronic pain populations.

Previously, a negative correlation has been found between somatosensory alpha power and perceived pain intensity for experimentally induced pain (Babiloni et al. 2006; Tu et al. 2016) and between frontal and somatosensory alpha power and chronic pain intensity (D. Camfferman et al. 2017). Ahn et al. (2019) also found that the increase of frontal and somatosensory alpha power by alpha tACS was associated with pain relief. In the present study, increasing alpha power with visual stimulation did not result in a significant reduction of pain intensity and unpleasantness. Moreover, the present study only found a non-significant negative correlation between standardized somatosensory alpha power (right-middle ROI) and the change in pain intensity (r = -.40; p = .082) and unpleasantness (r = -.41; p = .073) following alpha stimulation (sitting condition). As these correlations were only marginally significant and based on a relatively small sample (N = 20), no confident conclusions can be drawn from these findings. However, where this study only included brief periods of stimulation, Ahn et al. applied alpha tACS for 40 minutes. In an experimental pain setting with pain-free volunteers, Ecsy et al. (2018; 2017) achieved a significant reduction in pain ratings using 10 minutes of auditory and visual stimulation and Arendsen et al. (2018) applied alpha tACS for 15-20 minutes. This feasibility study focused primarily on the entrainment of alpha activity, where it has been shown that even very short periods of stimulation can entrain alpha oscillations (Herrmann 2001; Mathewson et al. 2012; Notbohm and Herrmann 2016). However, to also reduce chronic pain longer stimulation periods might be required. Moreover, this feasibility study included a relatively small group of participants, which introduces the possibility that the study is simply underpowered to find an effect of the stimulation on chronic pain. Further investigation with a larger sample size is needed to confirm whether a longer period of visual alpha stimulation leads to a significant reduction of chronic pain.

Further inspection of the individual changes in pain intensity and unpleasantness in response to the alpha stimulation showed that a wide variability in pain response was present (Figure 7, Table 5-8). Whereas some patients showed a reduction of several points on the 11-point NRS, others did not improve at all or even showed an increase of pain. Large variability in response is a problem for neurostimulation techniques in general. To improve the efficacy of neurostimulation interventions to manage chronic pain, it is important to take into account inter- and intra-individual factors such as cognitive, psychological, and neurophysiological state, and methodological factors that might contribute to this variability (Li, Uehara, and Hanakawa 2015; Fertonani and Miniussi 2017). In this study we did not identify a relationship between patient characteristics (as assessed with the questionnaires) and the pain response. However, larger sample sizes (e.g., 80-100) are likely needed for such analyses to be adequately powered for medium effect sizes. Another important source of variability in the effects of neurostimulation is brain-state dependency, i.e., the effect of neurostimulation depends on the timing of stimulation with respect to the underlying brain state. A number of studies have shown that applying neurostimulation in a brain-state dependent manner can enhance the modulation of corticospinal excitability (Kraus et al. 2016; Kaneko et al. 2014; Saito et al. 2013). Ultimately, taking into account these factors in the application of neurostimulation should lead to a more personalized and adaptive neuromodulatory therapy to reduce chronic pain.

Evidence shows that the efficacy of alpha entrainment depends on the distance between the stimulation frequency and the individual alpha peak frequency (IAF) (Herrmann et al. 2016; Notbohm, Kurths, and Herrmann 2016; Gulbinaite et al. 2017). Thus, tailoring the frequency of the visual alpha stimulation to each individual could improve the effect of alpha stimulation in patients with chronic pain. Moreover, recent studies have also explored the potential of combined stimulation. Anodal tDCS over the primary motor cortex (M1) combined with peripheral electrical stimulation led to an enhanced, long-lasting, and clinically important reduction in chronic pain (Schabrun et al. 2014; Hazime et al. 2017; Boggio et al. 2009). Together, these recent developments in the application of neurostimulation offer promising future directions for application of alpha stimulation to reduce chronic pain.

To successfully implement visual alpha stimulation to reduce chronic pain it is also important to better understand the relationship between alpha activity and chronic pain. So far, most studies have focused on the role of alpha activity in the perception of experimentally induced pain in pain-free individuals. Although there are some initial findings showing a negative correlation between frontal and somatosensory alpha power and chronic pain (Ahn et al. 2019; D. Camfferman et al. 2017), the functional role of alpha activity in the perception of chronic pain remains unclear. Experimental pain studies have demonstrated that the relationship between alpha activity and pain is influenced by attention (Hauck et al. 2015; May et al. 2012) and expectations about pain (Huneke et al. 2013; Arendsen, Hugh-Jones, and Lloyd 2018), and that pain expectations can influence the effect of neuro-stimulation on pain perception (Arendsen, Hugh-Jones, and Lloyd 2018). However, the relationship between attention, expectation, and alpha activity in a setting of chronic pain is little understood. It is important to better understand how these factors influence the relationship between alpha activity and chronic pain and the effectiveness of alpha stimulation to reduce chronic pain.

The present study showed that visual alpha stimulation offers a means to modulate alpha activity in patients with chronic pain in a lab-based environment. Whereas this is an important first step, further development is required to transform this lab-based application into a therapeutic technique that patients can use in their own home with therapeutic benefit. In a parallel study (Locke et al. 2020), a first qualitative assessment of a smartphone-based alpha entrainment technology was carried out. Individuals with chronic pain were asked about their experience with using the technology at home, using a virtual-reality headset for rhythmic visual stimulation and headphones for rhythmic auditory stimulation (binaural beats). The study provided initial support for the acceptability and usability of this smartphone-based technology as an affordable and accessible alternative to manage chronic pain. An important next step is to investigate the effectiveness of longer periods and multiple sessions of alpha stimulation to reduce chronic pain in the lab and at home, to translate these initial findings into a technology that can effectively reduce pain in a home-based setting.

To conclude, this study provides first evidence that visual stimulation at alpha frequency can be used to increase alpha power in patients with musculoskeletal pain. However, a brief 4-minute period of stimulation was not sufficient to reduce chronic pain. This study is a first step in the development of a novel neurostimulation approach to reducing chronic pain. Further study is warranted to investigate optimal dose and individual stimulation parameters (Krause and Cohen Kadosh 2014) to achieve significant pain relief. Together with the further development of a home-based neurostimulation platform, this could ultimately lead to the implementation of alpha stimulation as an affordable and accessible neurotherapy to manage chronic pain.

